# Bacterial protein interaction networks: connectivity is ruled by gene conservation, essentiality and function

**DOI:** 10.1101/2020.04.03.024265

**Authors:** Maddalena Dilucca, Giulio Cimini, Andrea Giansanti

## Abstract

Protein-protein interaction (PPI) networks are the backbone of all processes in living cells. In this work we relate conservation, essentiality and functional repertoire of a gene to the connectivity *k* of the corresponding protein in the PPI networks. Focusing on a set of 42 bacterial species with reasonably separated evolutionary trajectories, we investigate three issues: i) whether the distribution of connectivity values changes between PPI subnetworks of essential and nonessential genes; ii) how gene conservation, measured both by the evolutionary retention index (ERI) and by evolutionary pressures (evaluated through the ratio *K*_*a*_*/K*_*s*_ and ENC plots) is related to the the connectivity of the corresponding protein; iii) how PPI connectivities are modulated by evolutionary and functional relationships, as represented by the Clusters of Orthologous Proteins (COGs). We show that conservation, essentiality and functional specialization of genes control in a quite universal way the topology of the emerging bacterial PPI networks. Noteworthy, a structural transition in the network is observed such that, for connectivities *k* ≥ 40, bacterial PPI networks are mostly populated by genes that are conserved, essential and which, in most cases, belong to the COG cluster J, related to ribosomal functions and to the processing of genetic information.

## 1 Introduction

To operate biological activities in living cells, proteins work in association with other proteins, often assembled in large complexes. Hence, knowing the interactions of a protein is important to understand its cellular functions. Moreover, a comprehensive description of the stable and transient protein-protein interactions (PPIs) within a cell would facilitate the functional annotation of all gene products, and provide insight into the higher-order organisation of the proteome [12, 17]. Several methodologies have been developed to detect PPIs, and have been adapted to chart interactions at the proteome-wide scale. These methods, combining different technologies, experiments and computational analyses, generate PPI networks of sufficient reliability, enabling the assignment of several proteins to functional categories[31, 44].

Moreover, the statistical study of bacterial PPIs over several species (meta-interactomes) has brought important knowledge about protein functions and cellular processes [39, 8]. This work contributes to this line of research. Our aim is to shed light on the relationship between conservation, essentiality and functional annotation at the genetic level with connectivity patterns of PPI networks. We extend here our previous observations made on the PPI of *E.coli* which suggested a strong correlation between codon bias and the topology of PPI networks on the one hand, and between codon bias and gene conservation and essentiality on the other hand [11, 10]. It is worth, in the next two paragraphs, to make more precise what is usually meant by gene essentiality and gene conservation. Individual genes in the genome contribute differentially to the survival of an organism. According to their known functional profiles and based on experimental evidence, genes can be divided into two categories: essential and nonessential ones[16, 13]. Essential genes are not dispensable for the survival of an organism in the environment it lives in[13, 33]. Nonessential genes are instead those which are dispensable [28], being related to functions that can be silenced without compromising the survival of the organism. Naturally, each species has adapted to one or more evolving environments and, plausibly, genes that are essential for one species may be not essential for another one.

It has been argued many times that essential genes are more conserved than nonessential ones [20, 25, 29, 22, 1]. The term conservation” has, however, at least two meanings. On the one hand, a gene is conserved if orthologous copies of it are found in the genomes of many species, as measured by the Evolutionary Retention Index (ERI) [16, 5]. On the other hand, a gene is (evolutionarily) conserved when it is subject to a purifying, selective, evolutionary pressure, which disfavors mutations. This pressure can be measured, as usually it is, by *K*_*a*_*/K*_*s*_, the ratio of the number of non synonymous substitutions per non synonymous site to the number of synonymous substitutions per synonymous site. To measure the evolutionary pressures exerted either on low, intermediate and high connectivity proteins we use here both *K*_*a*_*/K*_*s*_ and the widely used ENC plots. In the second meaning a conserved gene is, in a nutshell, a slowly evolving gene [20, 19].

In this work we show that bacterial PPI networks display an interesting topological-functional transition, ruled by protein connectivity *k* and with a threshold between *k* = 40 and *k* = 50. Proteins with high PPI network connectivities (hubs) likely correspond to genes that are conserved and essential. Conversely, genes that correspond to hub proteins in the PPI network are likely to be essential, highly shared and subject to a more purifying evolutionary pressure, than the genes coding for proteins with a low *k*. Additionally, below the threshold the functional repertoire of proteins is heterogeneous, whereas, above the threshold there is a quite strict functional specialization.

## Materials and Methods

We consider a set of 42 bacterial genomes (that we have previously investigated in [10]), reported in Table 1. These genomes were chosen in order to have a reasonably large coverage of data concerning conservation, essentiality and selective pressure. To check the stability of the results shown in Figure 1 we have evaluated the connectivities of PPIs in an alternative set of bacterial species in Table 1. Nucleotide sequences were downloaded from the FTP server of the National Center for Biotechnology Information [4].

**Table 1.**
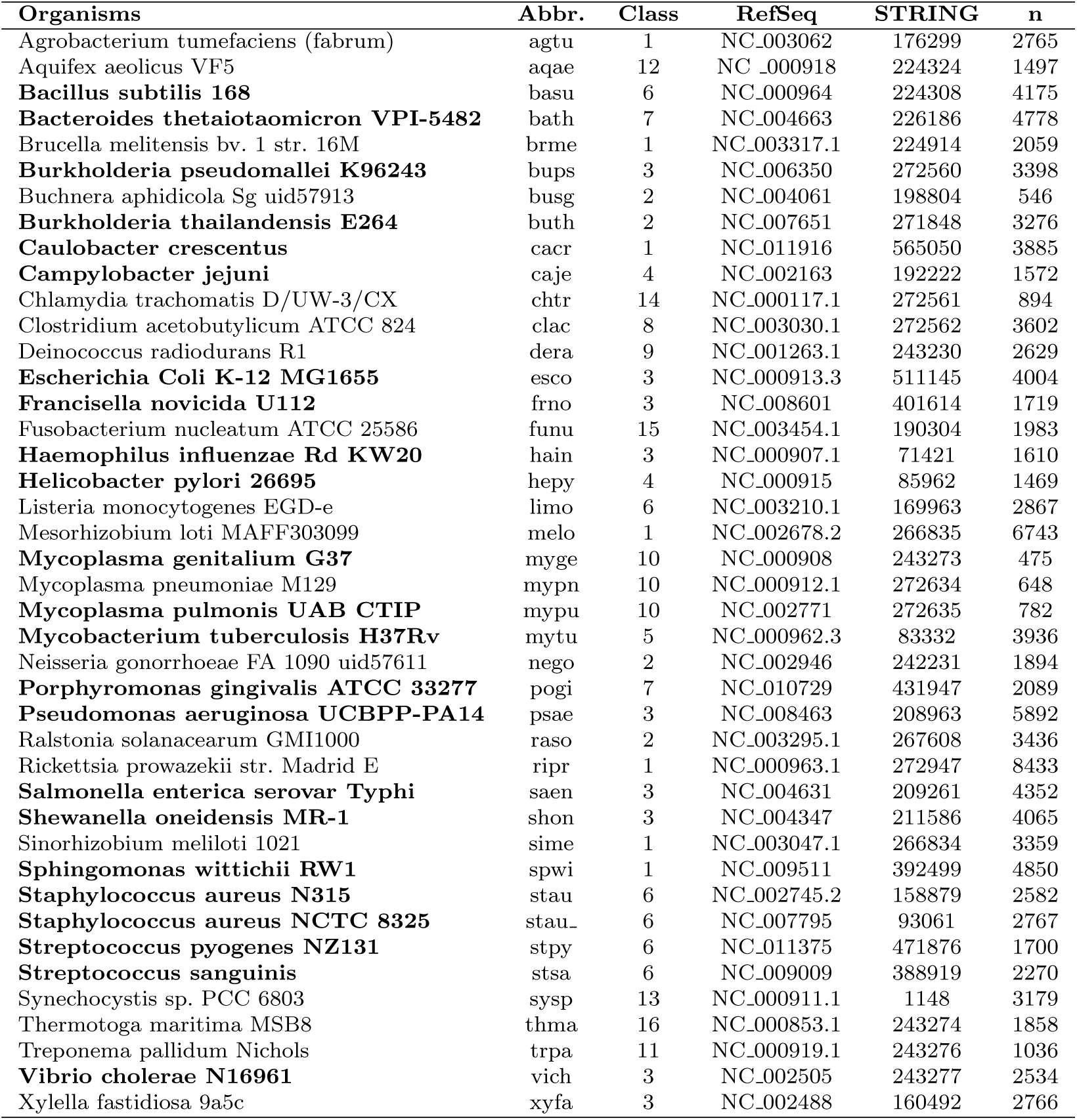
Summary of the selected bacterial dataset. Organism name, abbreviation, class, RefSeq, STRING code, size of genome (number of genes *n*). Genomes annotated in the Database of Essential Genes (DEG) are highlighted with bold fonts. Classes are:Alphaproteobacteria(1),Betaproteobacteria(2),Gammaproteobacteria(3),Epsilonproteobacteria(4),Actinobacteria(5),Bacilli(6),Bacteroidetes

**Fig. 1.**
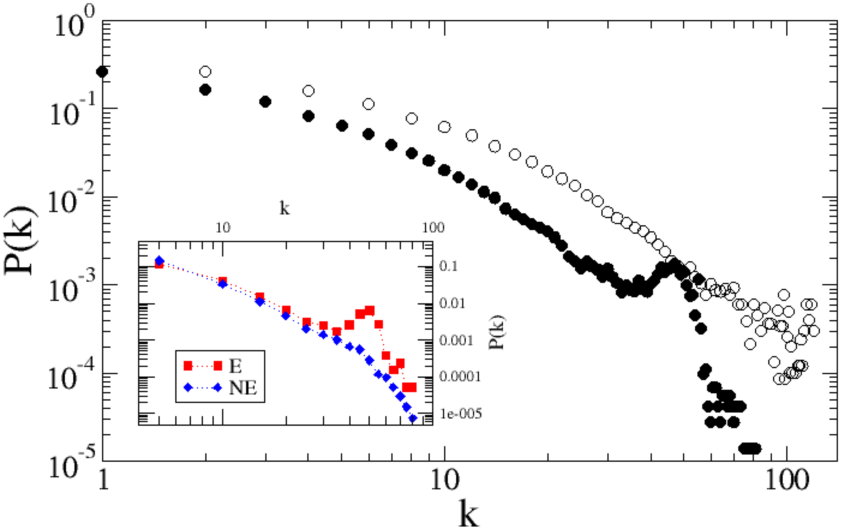
Probability distribution *P*(*k*) for the number of connections *k* of each protein, averaged over the bacterial species considered in Table 1 (full dots), compared with the degree distribution after removal of the proteins corresponding to genes in COG J, related to translational processes (empty dots). Inset: *P*(*k*) for essential (E) and nonessential (NE) genes, averaged over DEG-annotated genomes. Note that the average degree is higher for essential genes than for nonessential ones, and the two probability distributions are quite distinct. The region of the curve for low *k* can be well approximated by a power law [2].

### Gene Conservation

The Evolutionary Retention Index (ERI) [16] is a way of measuring the degree of conservation of a gene. In the present study the ERI of a gene is the fraction of genomes, among those reported in Table 1, that have at least an orthologous of the given gene. Then, as reminded in the Introduction, a low ERI value is related to a gene which is rather specific, common to a small number of genomes; whereas high ERI is characteristic of highly shared, putatively universal and essential genes.

We also make reference to another notion of gene conservation. Conserved genes are those which are subject to a purifying, conservative evolutionary pressure. To discriminate between genes subject to purifying selection and genes subject to positive selective Darwinian evolution, we use a classic but still widely used indicator, the ratio *K*_*a*_/*K*_*s*_ between the number of non synonymous substitutions per non synonymous site (*K*_*a*_) and the number of synonymous substitutions per synonymous site (*K*_*s*_) [19]. Conserved genes are characterized by *K*_*a*_*/K*_*s*_ *<* 1. We used *K*_*a*_*/K*_*s*_ estimates by Luo [29] that are based on the method by Nej and Gojobori[32].

### Gene Essentiality

We used the Database of Essential Genes (DEG, www.essentialgene.org) [29], which classifies a gene as either essential or nonessential on the basis of a combination of experimental evidence (null mutations or trasposons) and general functional considerations. DEG collects genomes from Bacteria, Archaea and Eukarya, with different degrees of coverage[48, 30]. Of the 42 bacterial genomes we consider, only 23 are covered—in toto or partially—by DEG, as indicated in Table 1.

### Protein-Protein Interaction Networks

PPIs are obtained from the STRING database (Known and Predicted Protein-Protein Interactions, https://string-db.org/)[41]. We have chosen STRING because of its quite large coverage of different bacterial species, useful to extend to multiple species the study we did in [11]. In STRING, each interaction is assigned with a confidence level or probability *w*, evaluated by comparing predictions obtained by different techniques [9, 34, 35] with a set of reference associations, namely the functional groups of KEGG (Kyoto Encyclopedia of Genes and Genomes)[26]. In this way, interactions with high *w* are likely to be true positives, whereas, a low *w* possibly corresponds to a false positive. As usually done in the literature, we consider only interactions with *w* ≥ 0.9 that allows for a fair balance between coverage and interaction reliability (see for instance the case of *E.coli* reported in [11]). We denote by *k* the *degree* (number of connections) associated to each proteins in each PPI network after the thresholding procedure. Note also that after applying the cut-off we are left, for each network, with a number of isolated proteins (with no connections) that grows as 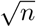 (where *n* is the number of proteins in the genome). These proteins are not considered in the network analysis and are regarded as stemming from statistical noise. In order to check the robustness of the bump in Figure 1 we have also considered the degree distributions relaxing the cut-off values to *w* = 0.0, 0.5 and 0.7; see Figures 1, 2 and 3 of Supplementary Information.

**Fig. 2.**
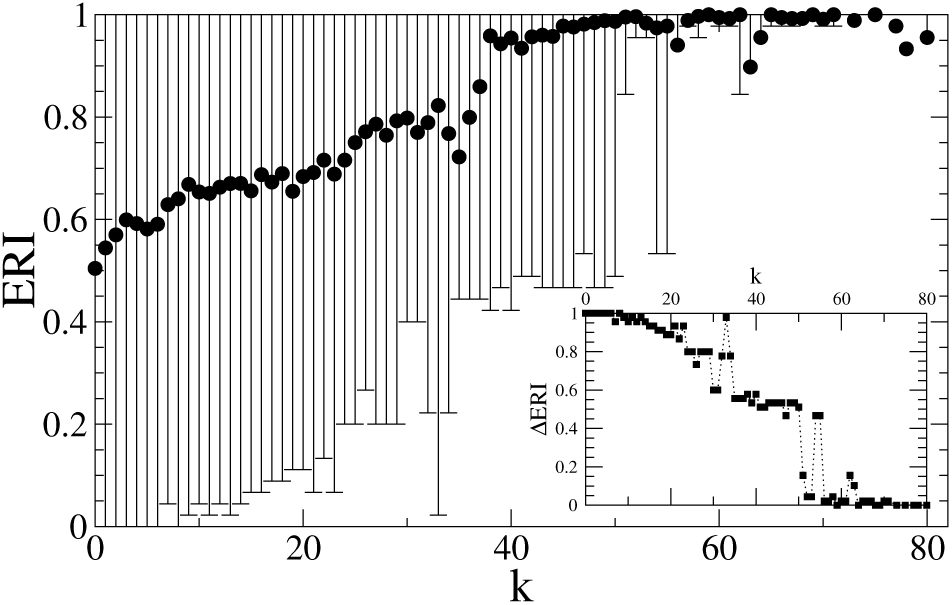
Average ERI values of bacterial genes as a function of the degrees *k* of the corresponding proteins, for all the considered genomes. Error bars are standard deviations of ERI values associated to a given *k* value. Inset: amplitude of the error bar (*Δ*ERI) as a function of *k*.

**Fig. 3.**
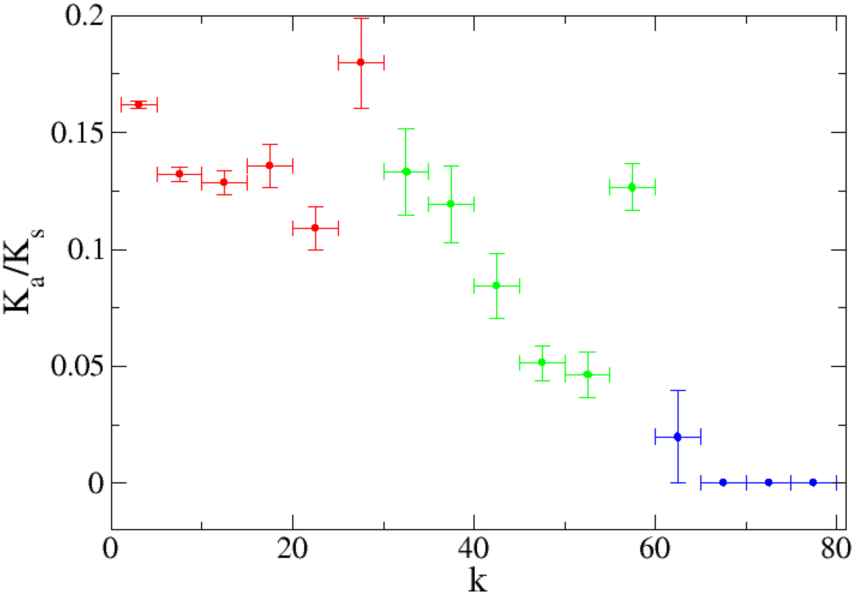
Dependence of purifying selective pressure (*K*_*a*_*/K*_*s*_) in groups of genes corresponding to proteins with different connectivity degrees *k*.

### K_a_/K_s_

*K*_*a*_/*K*_*s*_ is the ratio of nonsynonymous substitutions per nonsynonymous site (*K*_*a*_) to the number of synonymous substitutions per synonymous site (*K*_*s*_) [19]. This parameter is widely accepted as a straightforward and effective way of separating genes subject to purifying evolutionary selection (*K*_*a*_*/K*_*s*_ *<* 1) from genes subject to positive selective Darwinian evolution (*K*_*a*_*/K*_*s*_ *>* 1). There are different methods to evaluate this ratio, though the alternative approaches are quite consistent among themselves. For the sake of comparison, we have used here the *K*_*a*_*/K*_*s*_ estimates by Luo et al. [29] which are based on the Nej and Gojobori method [32]. Note that each genome has a specific average level of *K*_*a*_/*K*_*s*_. In Figure 3 average values of *K*_*a*_*/K*_*s*_ are shown for low, intermediate, high connectivity bins of genes.

### ENC plot

The ENC-plot is a well known tool to investigate the patterns of synonymous codon usage in which the *ENC* values are plotted against *GC*_3_ values when codon usage is dominated by the mutational bias, the formula of expected *ENC* values is given by:

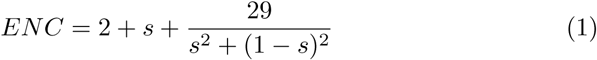

*s* represents the value of *GC*_3_ [46]. When the corresponding points fall near the expected neutral curve, mutations that enforce the typical mutational bias of the species are the main factor affecting the observed codon diversity. Whereas when the corresponding points fall considerably below the expected curve, the observed CUB is mainly affected by natural selection. To quantitatively represent the balance between mutational bias and selective natural pressure we parametrize the ENC formula, to be used in non-linear fits to the experimental data:

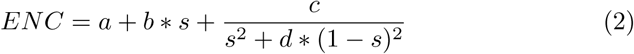

### Clusters of orthologous proteins

We use the functional annotation given in the database of orthologous groups of proteins (COGs) from Koonin’s group, available at *http://ncbi.nlm.nih.gov/COG/*[42, 15]. We consider 15 functional COG categories (see Table 2), excluding the generic categories R and S for which functional annotation is too general or missing.

**Table 2.** Functional classification of COG clusters.

| COG ID | Functional classification |
| --- | --- |
|  | *INFORMATION STORAGE AND PROCESSING* |
| J | Translation, ribosomal structure and biogenesis |
| K | Transcription |
| L | Replication, recombination and repair |
|  | *CELLULAR PROCESSES AND SIGNALING* |
| D | Cell cycle control, cell division, chromosome partitioning |
| T | Signal transduction mechanisms |
| M | Cell wall/membrane/envelope biogenesis |
| N | Cell motility |
| O | Post-translational modification, protein turnover, chaperones |
|  | *METABOLISM* |
| C | Energy production and conversion |
| G | Carbohydrate transport and metabolism |
| E | Amino acid transport and metabolism |
| F | Nucleotide transport and metabolism |
| H | Coenzyme transport and metabolism |
| I | Lipid transport and metabolism |
| P | Inorganic ion transport and metabolism |

## Results and discussion

### Degree distribution of PPI networks

We start by studying the degree distributions *P* (*k*) observed in bacterial PPIs. We first recall that such a distribution was found to be scale-free in *E.coli* [7, 24, 36, 11], meaning that the corresponding PPI network features a large number of poorly connected proteins, and a relatively small number of highly connected hubs. In order to assess the generality of this observation, we compute *P* (*k*) for each genome in Table 1 (plots are reported in Figures S4-S5 of the Supplementary Information). Note that, despite that PPIs of different bacteria have different sizes and densities, their average connectivity and the support of their *P* (*k*) are very similar (as shown in Figure S1 of the Supplementary Information). Thus, we can superpose all the considered bacterial degree distributions without the need to normalise the support of each *P* (*k*). When doing so, we observe two distinct regimes (see Figure 1). For high values of *k*, the distribution is scale-free: *P* (*k*) ∝ *k*^−*γ*^. This scaling behaviour is consistent with previous studies on the genomes of yeast, worms and flies [18] and on co-conserved PPIs in some bacteria[27]. For higher values of *k*, however, the distribution deviates from a power law, and a bump with a Gaussian-like shape emerges. Interestingly, this feature is almost undetectable in the graph relative to a single species (such as, e.g., emph E.coli, see Figure 2 in Supplementary Information of Dilucca et al. [11]), but clearly emerges when the statistics is enriched by adding together the graphs relative to several species. This feature, emerging for *k* ≥ 40 is reasonably due to the contribution of proteins belonging to complexes [47]. In particular, if one re-calculates the degree distribution of a data set in which the ribosomal proteins are removed the bump is not present (see figure1, empty dots). Moreover, if we consider the separate contribution of essential and nonessential genes to the *P* (*k*) (for DEG-annotated genomes), we see that the superposed peak is present only in the degree distribution of essential genes. Moreover, the degree distributions for essential and nonessential genes are well separated and the average degree is systematically higher for essential genes than for nonessential ones—consistently with previous findings [18].

A critical remark is in order at this point. One could ask how many species are needed to obtain the bump observed in figure 1 as a stable feature? To check this point we have evaluated the degree distributions obtained by gradually averaging over an increasing number of species, taken in an arbitrary order.

As shown in figure S2 it is sufficient to average the *P* (*k*) over no more than 10 species to have the bump consistently emerge and stabilise as a self-averaging feature, that is mainly associated to the complex of ribosomal proteins. Moreover, we have checked that the observed bump is still present averaging the degree distributions in the PPIs of 42 alternative bacterial species (see figure S3). Then, we can conclude that it is sufficient to average over just a few species to let the the bump emerge as a general feature.

### PPI connectivity and gene conservation

We now investigate whether the connectivity *k* of a protein in a PPI network drives a transition in the degree of conservation (as measured by ERI) of the corresponding genes. Figure 2 displays the average value and the spread of ERI in genes relative to bins of proteins that are iso-connected in the PPIs of different species. As a general feature we observe that, on the average, the genes of highly connected proteins are highly conserved among the bacterial species we consider, that constitute a reasonably wide sample of different evolutionary adaptations. The same figure 2 shows that if *k* ≤ 50 then the ERI highly fluctuates between different samples of proteins with the same *k*, in different species. For high connectivities (above *k* = 50), the ERI is close to 1, with a drastic drop in the fluctuation (as shown in the inset). This observation points to the existence, in each bacterial PPI, of an almost-invariant structure of conserved hubs, sustained by highly conserved genes. We have checked that a connectivity of 40 acts as a lower bound, then we can conclude, as a rule of thumb, that a protein with connectivity degree of 40 or more is likely to be coded by a gene shared by at least 80% of the species in a generic pool of bacteria. At the moment is hard to figure out deep and general biophysical facts that could give a universal character to the threshold we are observing here, in the form of a new biological law. Let us just propose, as an heuristic observation, the existence of a characteristic, almost-critical value of connectivity to be set close to 50.

### Evolutionary pressure and PPI connectivity

We then look at the evolutionary pressure exerted on genes whose proteins have different connectivities. The graph in figure 3 shows the ratio *K*_*a*_*/K*_*s*_ for groups of genes binned by the connectivity *k* of the corresponding proteins, for all the 42 bacterial species in table 1. As is well known this ratio *K*_*a*_*/K*_*s*_ provides a straightforward indication of the balance between a positive driving *darwinian selection* (when the numerator prevails) and a *purifying*, stabilizing selection (acting against change in genes for which the denominator prevails).

We see that the more connected proteins correspond to genes which are subject to an increasing purifying evolutionary pressure. Indeed, (*K*_*a*_*/K*_*s*_) is less than 1 in all bins of connectivity and systematically decreases, as a function of *k*, down to zero. A decreasing ratio generally indicates an increasing role of the purifying conservative selection in the corresponding set of genes.

To add evidence to this observation we have also considered ENC plots for sets of genes binned by the connectivities of the corresponding proteins. Interestingly, the data in figure 4 are fully consistent with those in figure 3. In the ENC plots, the points associated to low connectivity proteins (green) are closer to the so called Wright’s profiles (represented as black solid lines) than those associated to proteins with intermediate and high connectivities (red and blue lines). Figure 5 stresses this observation in a more quantitative way by showing that in the ENC plots the average distance from Wright’s profile monotonously increases with *k*, Overall, the above results clearly indicate that codon bias and GC content of high connectivity genes are more under selective darwinian pressure than genes coding for low-connectivity proteins, in which the rate of accepted mutations is mainly ruled by mutational bias.

**Fig. 4.**
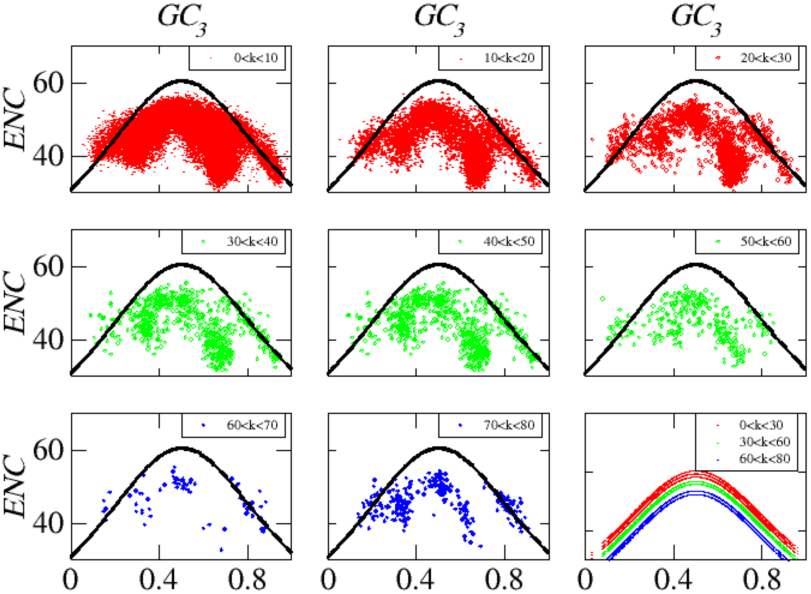
ENC plots for three groups of genes corresponding to proteins with different degree connectivities. In each panel the solid black lines are plots of Wright’s theoretical curve (equation 1) which correlates effective number of codons with *GC*_3_ in the case of pure mutational bias (no selective pressure). Coherently with figure 3 the case of low connectivities are shown in red, intermediate in green and high connectivities in blue. In the bottom-right panel dashed linear non-linear fits of Wright’s theoretical shapes to the experimental data. The fitted curves are shown together in the bottom right panel. For the sake of completeness the best fit parameters are reported in table 5.

**Fig. 5.**
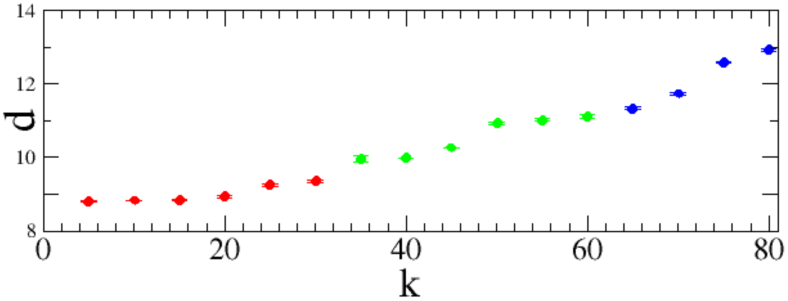
Average distance *d* from Wright’s curve as a function of *k*. The distance from the curve increases with *k*.

### PPI and Essentiality

To further investigate the relationship between gene essentiality and protein connectivities, we consider DEG-annotated genomes and classify interactions between proteins (links) making references to the essentiality of the corresponding genes. We distinguish three sets of links: *ee* (linking proteins from two essential genes), *ēē* (from two nonessential genes) and *eē* (from an essential gene and a nonessential one). We then compute the *density* of these sets of links respectively as:

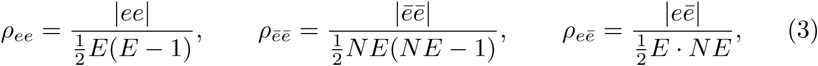

where *E* and *NE* denote the number of essential and nonessential genes, respectively (self-connections are excluded in our analysis). Such densities are then compared with the overall density of the network—restricted to genes classified as either essential or nonessential:

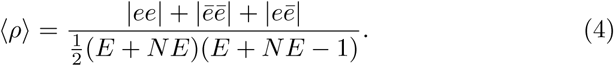

**Table 3.**
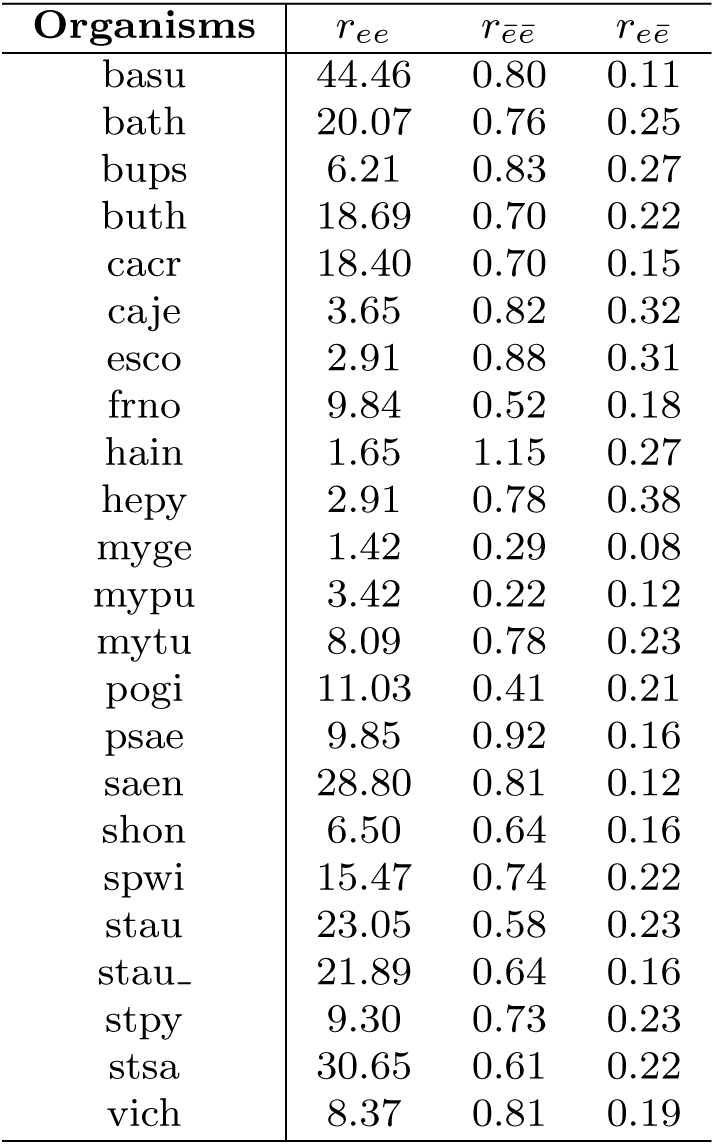
Relative density values *r* for PPI subnetworks between essential genes (*r*_*ee*_), between nonessential genes (*r*_*ēē*_) and between essential and nonessential genes (*r*_*eē*_), for each DEG-annotated bacterial genome.

**Table 4.**
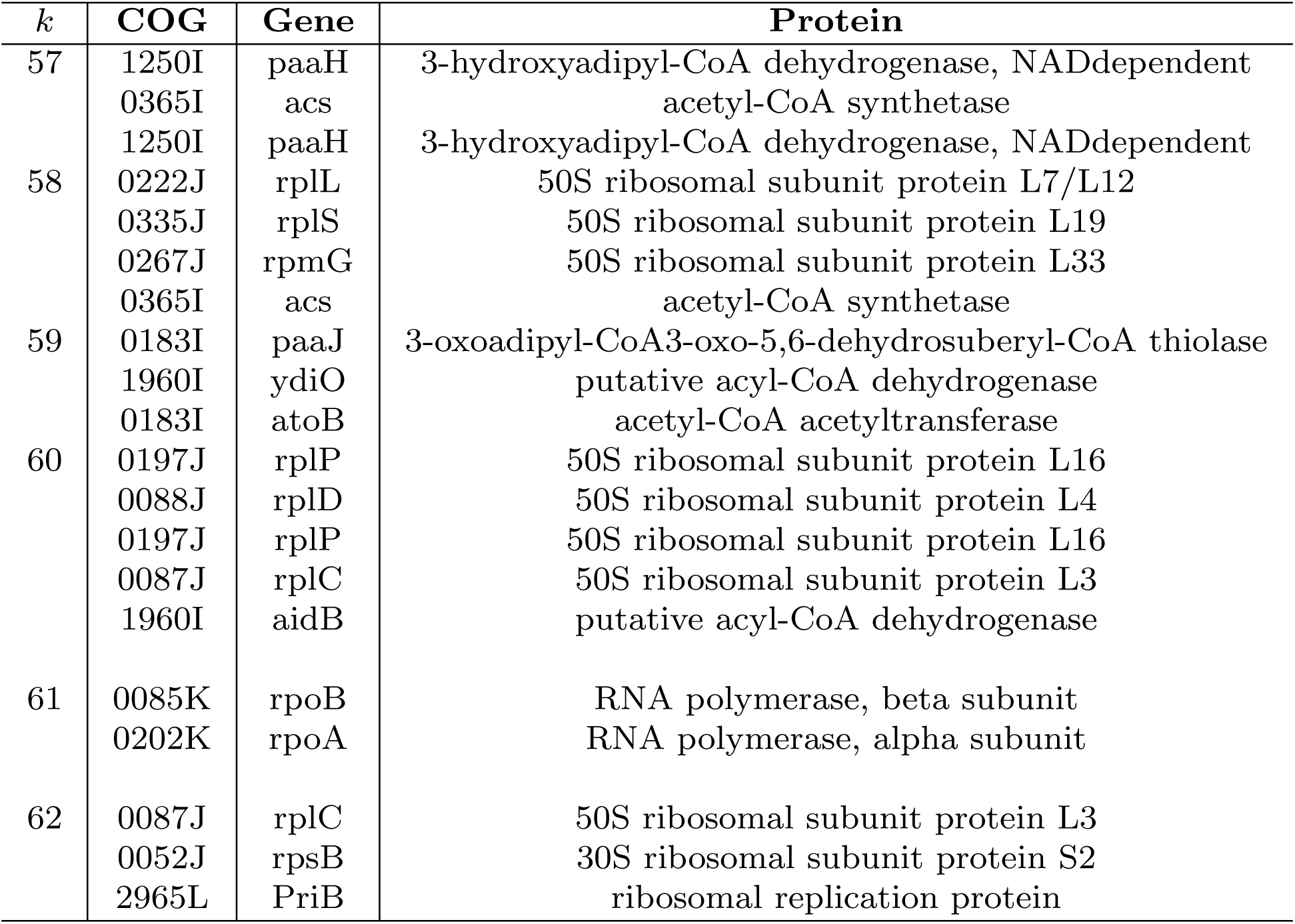
Specific hubs. In this table we detail which proteins populate the few bins of connectivity around *k* = 60 in figure 6.

**Table 5.** Best fit values of the parameters in equation 1 and correlation coefficients for different connectivity data, shown in figure 4.

| $k$ | $a$ | $b$ | $c$ | $d$ | $R^2$ |
| --- | --- | --- | --- | --- | --- |
| [0 – 10] | 40,561 | -10,338 | 5,555 | 1,052 | 0,617 |
| [10 – 20] | 23,774 | 3,890 | 8,583 | 0,626 | 0,590 |
| [20 – 30] | 20,280 | 8,287 | 8,276 | 0,507 | 0,540 |
| [30 – 40] | 18,296 | 10,685 | 8,334 | 0,190 | 0,790 |
| [40 – 50] | 18,548 | 10,372 | 8,326 | 0,495 | 0,589 |
| [50 – 60] | 25,868 | 2,508 | 8,038 | 0,650 | 0,758 |
| [60 – 70] | 29,344 | -0,756 | 10,224 | 0,977 | 0,870 |
| [70 – 80] | 30,507 | 3,438 | 6,990 | 0,811 | 0,874 |

We use the ratios *r*_*ee*_ = *ρ*_*ee*_*/* ⟨*ρ*⟩, *r*_*ēē*_ = *ρ*_*ēē*_*/* ⟨*ρ*⟩ and *r*_*eē*_ = *ρ*_*eē*_ ⟨*ρ*⟩ to assess the level of connectivity of the subnetworks with respect to the overall connectivity. Table 3 shows that subnetworks of essential genes are far denser than the overall networks, and that, in general, essential and nonessential genes tend to form network components that are weakly interconnected. This happens because many essential genes encode for ribosomal proteins, which in turn are localized in the ribosome so that they have a major probability to interact [3]. Figures S4-S5 of the Supplementary Information display such network features for each individual species.

### PPI connectivity and functional specialization

For each PPI network, we define the conditional probability that a protein with degree *k* belongs to a given COG as:

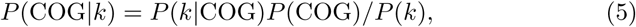

where *P* (*k*) is the degree distribution in the PPI network, *P* (COG) is the frequency of that COG in the proteome, and *P* (*k*|COG) is the degree distribution restricted to that COGs. Figure 6 shows the COG spectrum as a function of *k* over all bacteria species considered. Interestingly, we again note a marked transition. Below *k* ≃ 40 the COG spectrum is quite heterogeneous: genes corresponding to proteins with low connectivity are spread over several COGs which correspond to different functions (see Table 2).

**Fig. 6.**
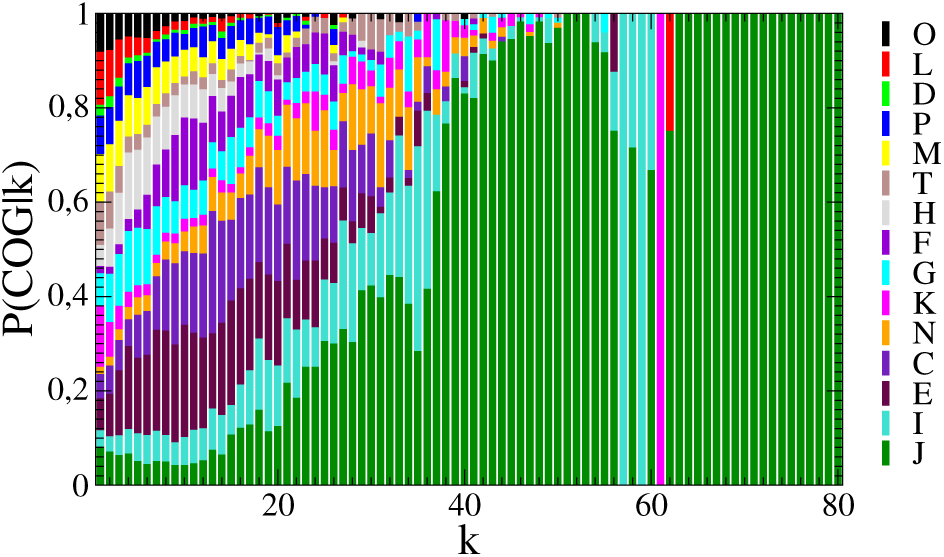
Probability distribution *P* (COG |*k*) of belonging to a given COG for proteins with degree *k*, over all considered genomes. Proteins with low connectivity have a very heterogeneous COG composition, whereas, those with high *k* basically belong only to COG J.

The transition shows that proteins with more than 40 interactions are likely to be coded by genes belonging to COG J, which encompasses translation processes and ribosomal functions. There are yet a handful of outliers, hubs with connectivities between 57 and 62, that belong to COG I (related to lipid transport and metabolism) and K and L (which, together with J, define the functional class of information storage and processing). The list of these outliers is reported in Table 4. But, which are the genes of COG J that drive the transition? In the next Figures 7 and 8 we are able to show which genes are the main characters in the transition. We investigated then the connectivities of the highly conserved (shared by all the species in Table1) genes belonging to COG J and whose proteins have connectivities bigger than 40. These highly shared genes corresponding to cores of highly connected ribosomal proteins are listed in Table 6. In the heatmap of Figure 7 we sort each gene in the COG J in order of ascending degree, species by species, and we see there is a core of genes (in red, lower left sector) that correspond to highly connected proteins, which are also highly shared (ERI=1, see Table 6) among all the species we considered. It is quite clear that in the heat map of Figure 7 the 42 species in this study can be splitted into at least four groups. Each group of species is characterized by different connectivities of their conserved ribosomal proteins. In other words, this observation suggests that the abrupt transition shown in Figure 6 is driven by a subset of COG J genes which are listed in Table 6 corresponds detailed connectivity of highly conserved ribosomal proteins. In order to show how conservation of the gene and connectivity of the protein are correlated we have built the connectivity map of the proteins corresponding to the COG J genes (Figure 8). The map in Figure 8 shows in a color code how much is conserved a given interaction (e.g., the red color corresponds to links in the PPI that are conserved among 37 species over 42). It is evident in this map that the set of highly conserved COG J genes of Table 6 corresponds to the core of highly connected ribosomal proteins, which are approximately invariant among several species.

**Table 6.** Genes belonging to COG J with average degree bigger than 40 (see Figures 7 and 8). All these genes are conserved, common to all species (ERI=1), and drive the transition shown in Figure 6

| COG | Genes name | <k> |
| --- | --- | --- |
| COG0097J | 50S ribosomal protein L6 | 60.24 |
| COG0087J | 50S ribosomal protein L3 | 60.19 |
| COG0197J | 50S ribosomal protein L16 | 60.19 |
| COG0090J | 50S ribosomal protein L2 | 60.14 |
| COG0080J | 50S ribosomal protein L11 | 60.12 |
| COG0088J | 50S ribosomal protein L4 | 60.12 |
| COG0081J | 50S ribosomal protein L1 | 58.19 |
| COG0089J | 50S ribosomal protein L23 | 57.88 |
| COG0102J | 50S ribosomal protein L13 | 57.45 |
| COG0094J | 50S ribosomal protein L5 | 57.21 |
| COG0092J | 30S ribosomal protein S3 | 57.12 |
| COG0098J | 30S ribosomal protein S5 | 57.10 |
| COG0093J | 50S ribosomal protein L14 | 57.00 |
| COG0091J | 50S ribosomal protein L22 | 56.24 |
| COG0049J | 30S ribosomal protein S7 | 55.31 |
| COG0051J | 30S ribosomal protein S10 | 55.24 |
| COG0200J | 50S ribosomal protein L15 | 55.12 |
| COG0256J | 50S ribosomal protein L18 | 54.86 |
| COG0203J | 50S ribosomal protein L17 | 54.43 |
| COG0244J | 50S ribosomal Protein L10 | 54.19 |
| COG0100J | 30S ribosomal protein S11 | 53.76 |
| COG0522J | 30S ribosomal protein S4 | 53.43 |
| COG0096J | 30S ribosomal protein S8 | 53.10 |
| COG0099J | 30S ribosomal protein S13 | 52.88 |
| COG0048J | 30S ribosomal protein S12 | 52.14 |
| COG0198J | 50S ribosomal protein L24 | 50.83 |
| COG0185J | 30S ribosomal protein S19 | 50.52 |
| COG0199J | 30S ribosomal protein S14 | 50.45 |
| COG0103J | 30S ribosomal protein S9 | 49.45 |
| COG0480J | tetracycline resistance protein. tetM | 47.90 |
| COG0052J | 30S ribosomal protein S2 | 47.69 |
| COG0184J | 30S ribosomal protein S15 | 45.95 |
| COG0186J | 30S ribosomal protein S17 | 44.60 |
| COG0255J | 50S ribosomal protein L29 | 43.95 |
| COG0222J | 50S ribosomal protein L7/L12 | 42.43 |
| COG1841J | 50S ribosomal protein L30 | 40.71 |

**Fig. 7.**
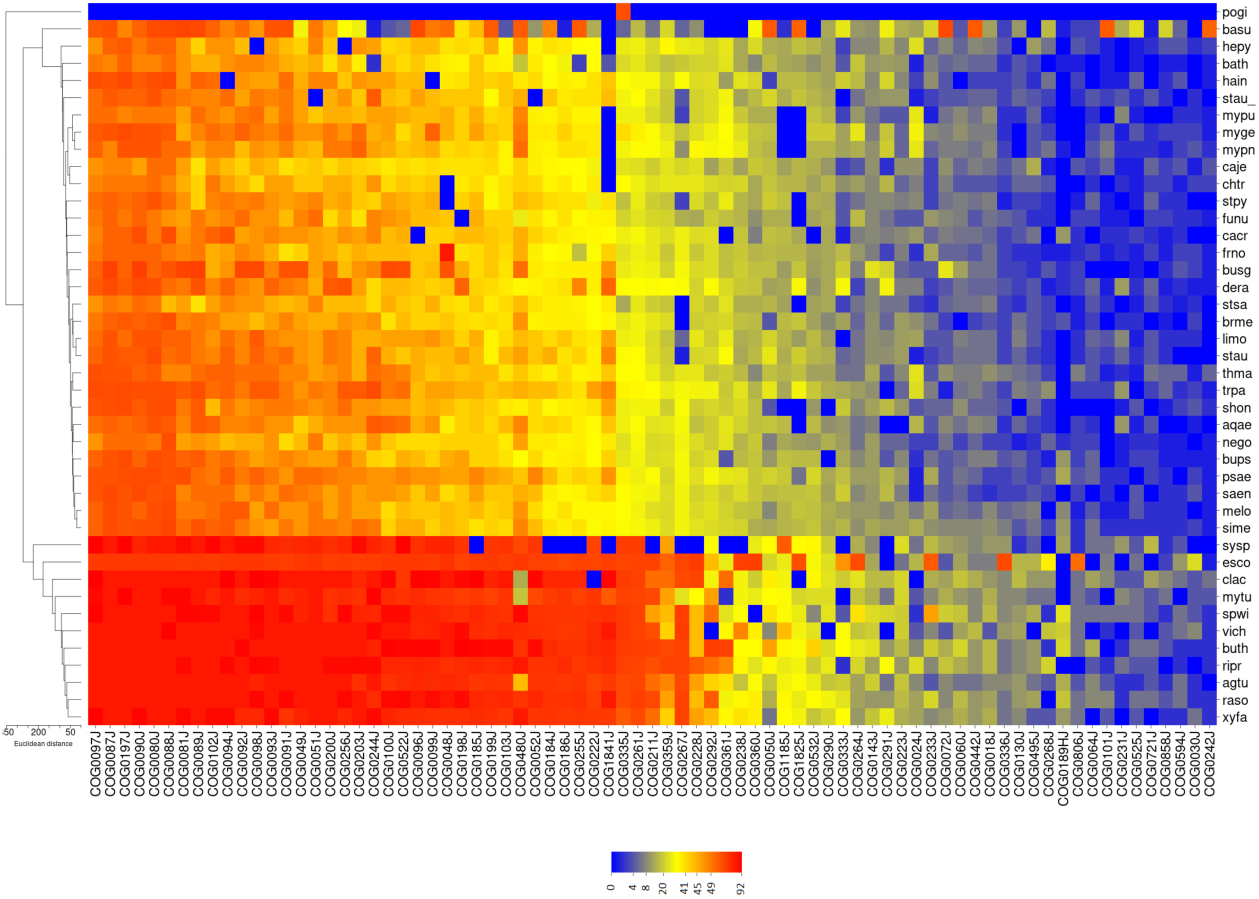
Heat map of degree of genes class COG J for each species. Genes are sorted by averaging degree. We note that genes with average degree major than 40 are conserved for all species. Details of these genes are in Table 6.

**Fig. 8.**
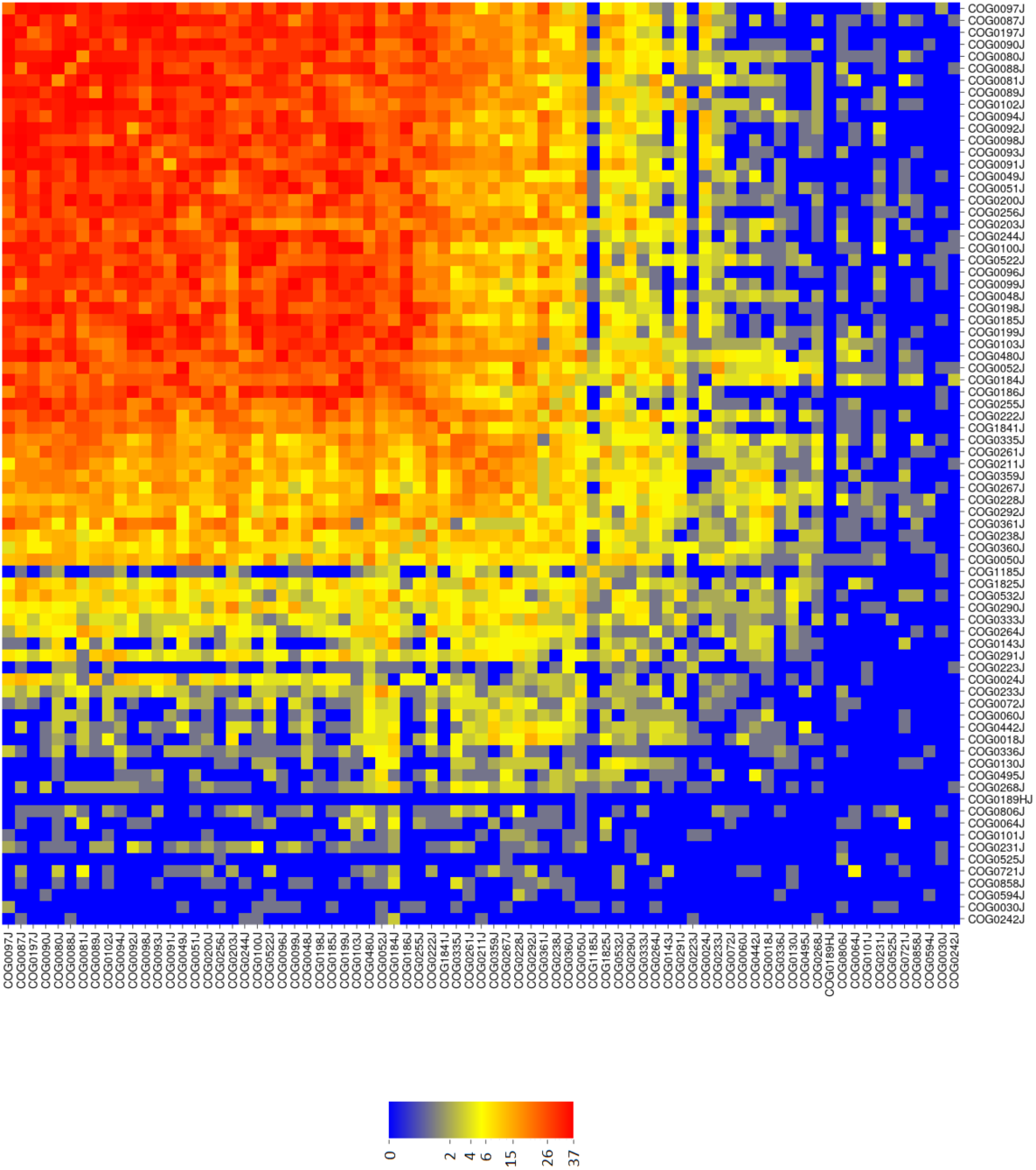
Connectivity map of the proteins coded by COG J genes. The color code measures the degree of conservation of interactions between proteins among the various species. When the link is present in all bacteria the number of conservation is 42, whereas when the link is not present the number is 0.

## Conclusions

Topological analysis of biological networks, such as protein-protein interaction or metabolic networks, has demonstrated that structural features of network subgraphs are correlated with biological functions [40, 37]. For instance, it was shown that highly connected patterns of proteins in a PPI are fundamental to cell viability [23]. In this work we have shown the existence of a topological-functional transition in bacterial species, ruled by the connectivity of proteins in the PPI networks. The threshold in *k* of the transition is located between *k* = 40 and *k* = 50. Proteins that have connectivities above the threshold are mostly encoded by genes that are conserved (as measured both by ERI and *K*_*a*_*/K*_*s*_) and essential. Moreover the functional repertoire above the threshold focuses mainly on the COG J (Translation, ribosomal structure and biogenesis), with just a few interesting hubs belonging to COGs I (Lipid transport and metabolism), K (Transcription) and L (Replication, recombination and repair).

Indeed, the PPI network of each bacterial species is characterized by a highly connected core of conserved ribosomal proteins, the components of multi-subunit complexes whose corresponding genes are mostly essential [7, 27] and code for supra-molecular complexes, that pile up in the bump we have observed for the degree distribution (figure1). Hence, what we are seeing here is essentially the ribosome, and related protein complexes such as RNA Polymerase. Indeed, the ribosome is the only molecular machine in bacteria in which a given protein could legitimately have 40 or more protein binding partners, with the help of rRNA mediating interactions [14].

We believe that the observations we have presented here can have implications both for the prediction of gene essentiality, based on the knowledge of PPI networks, and for the prediction of interactions between proteins, based on genetic information[21, 45]. It is interesting to note that our results are consistent with a previous study based on inferred bacterial co-conserved networks based on phylogenetic profiles [27]. The coupled flows of information from the genetic level up to the proteomic level and vice-versa should be further systematically investigated, taking into account archaeal and prokaryotic genomes in the search for emerging multi-layer structures that could offer basic theoretical grounds for clinical and systemic applications, for instance related to antimicrobial resistances [43, 49, 38, 6].

