## Supplemental file pdf for "Bacterial protein interaction networks: connectivity is ruled by gene conservation, essentiality and function"

### SUPPLEMENTARY INFORMATION

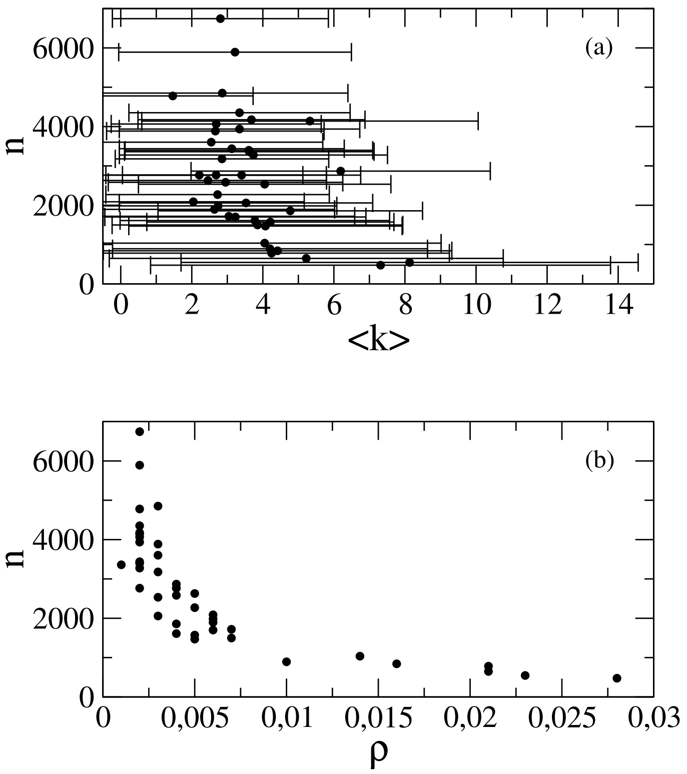


FIG. S1. Relation between genome size $n$ and average degree $\langle k \rangle \pm \sigma_k$ (upper panel) and density $\rho$ (bottom panel) of the corresponding PPI network for the set of bacterial species reported in Table 1 of the main text.

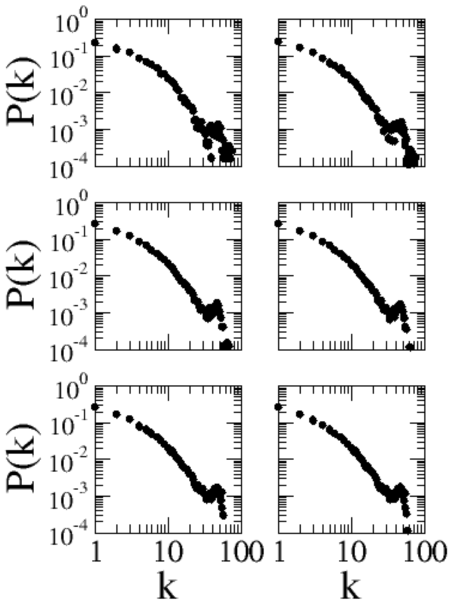


FIG. S2. Degree distributions $P(k)$ obtained by averaging over an increasing number of species considered in Table 5 of the main text, taken in an arbitrary order. From top left in the inset averaging over 5,10,15,20,25,30,35 and 42 species. Clearly, it is sufficient to average the $P(k)$ over no more than 15 species (first row) to have the bump consistently emerge and stabilise as a self-averaging feature, that is mainly associated to the complex of ribosomal proteins.

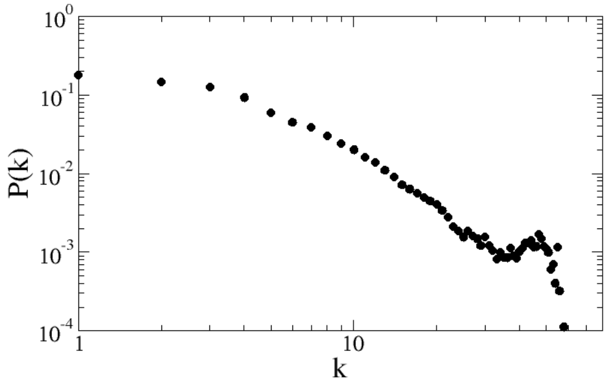


FIG. S3. Probability distribution $P(k)$ for the number of connections $k$ of each protein, avereged over the 42 alternative bacterial species listed here in the supplementary Table 1.

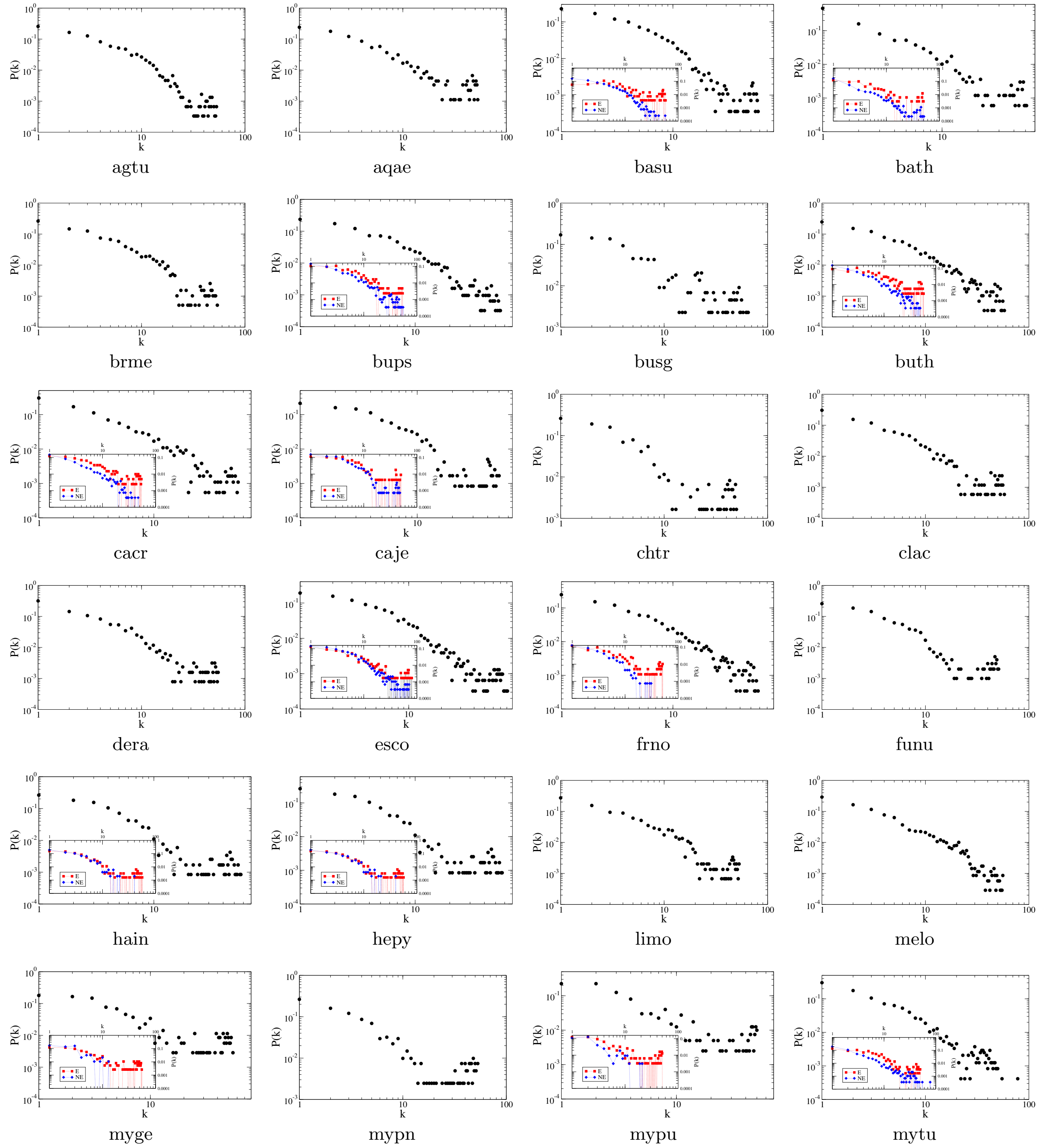


FIG. S4. Degree distribution $P(k)$ (part 1) of the PPI networks for the bacterial species reported in Table 1 of the main text. For DEG-annotated genomes, the inset shows the contribution of essential (red) and nonessential (blue) genes.

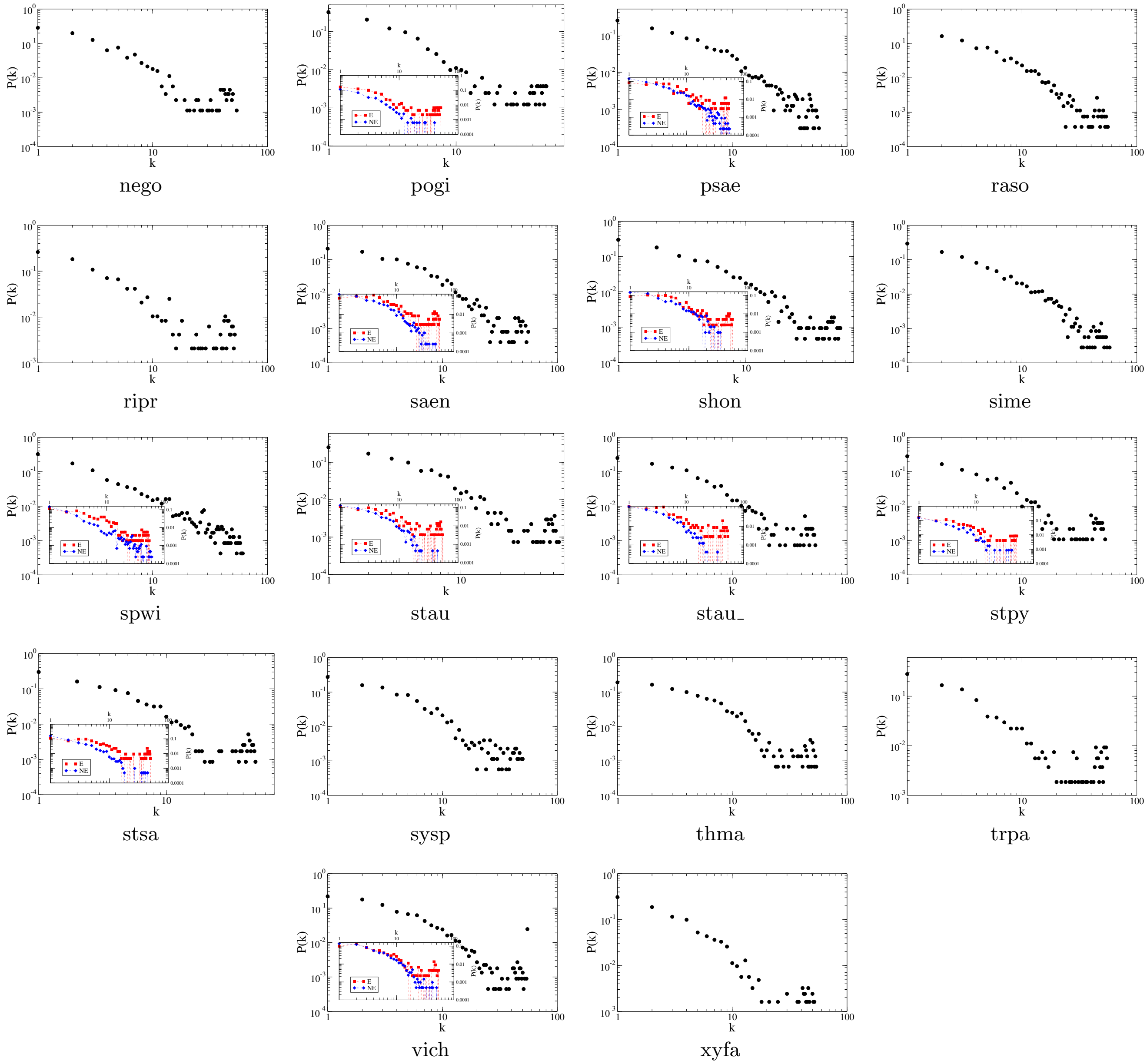


FIG. S5. Degree distribution $P(k)$ (part 2) of the PPI networks for the bacterial species reported in Table 1 of the main text. For DEG-annotated genomes, the inset shows the contribution of essential (red) and nonessential (blue) genes.

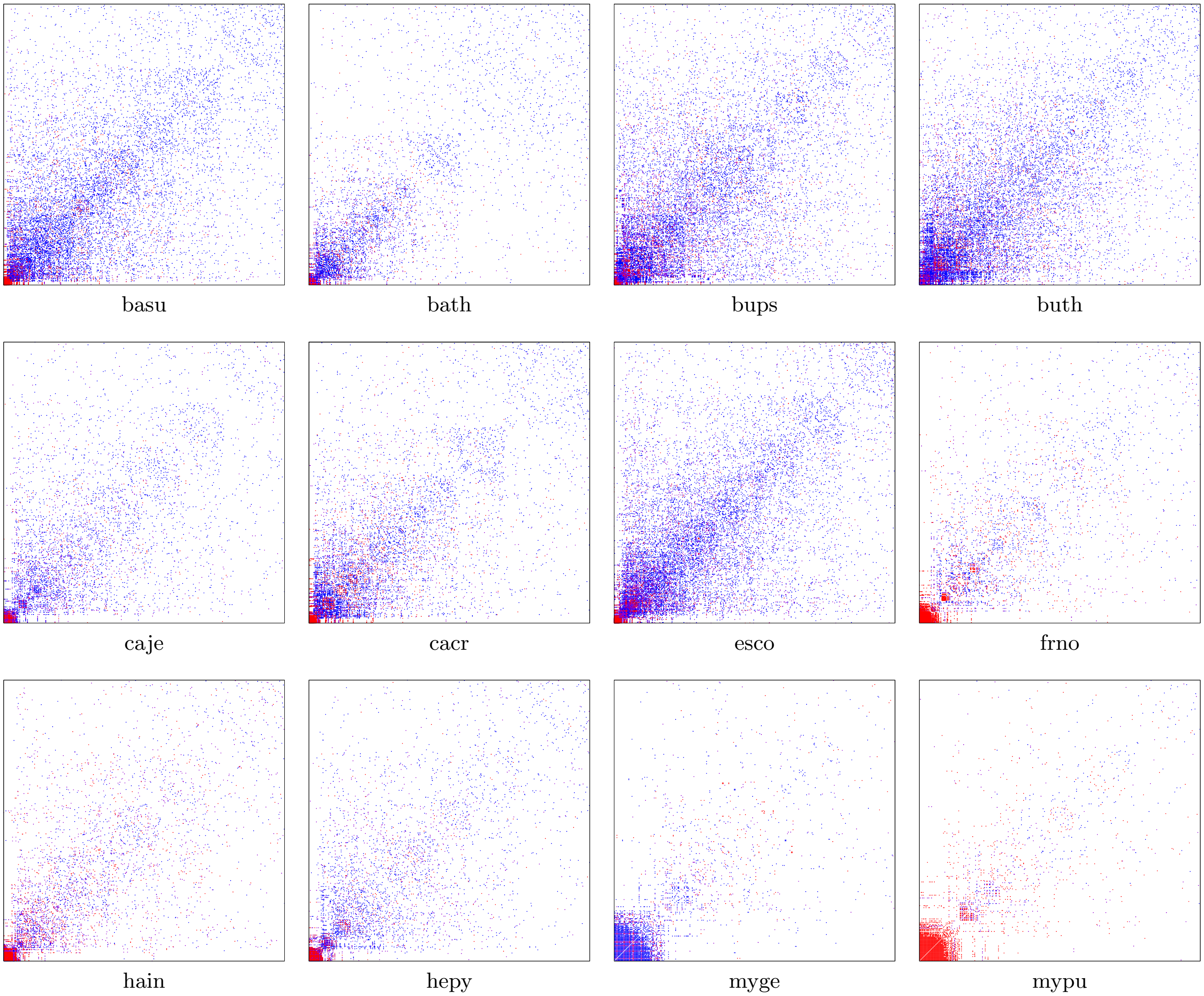


FIG. S6. Adjacency matrices (part 1) of the PPI networks for the DEG-annotated bacterial species reported in Table 1 of the main text. In each matrix, genes are ordered according to the degree of the corresponding protein in the network, in descending order from left to right and from bottom to top. Links among essential genes correspond to red-colored dots, those among nonessential (and non-annotated) genes to blue-colored dots, and those between essential and nonessential (plus non-annotated) genes to a violet-colored dot.

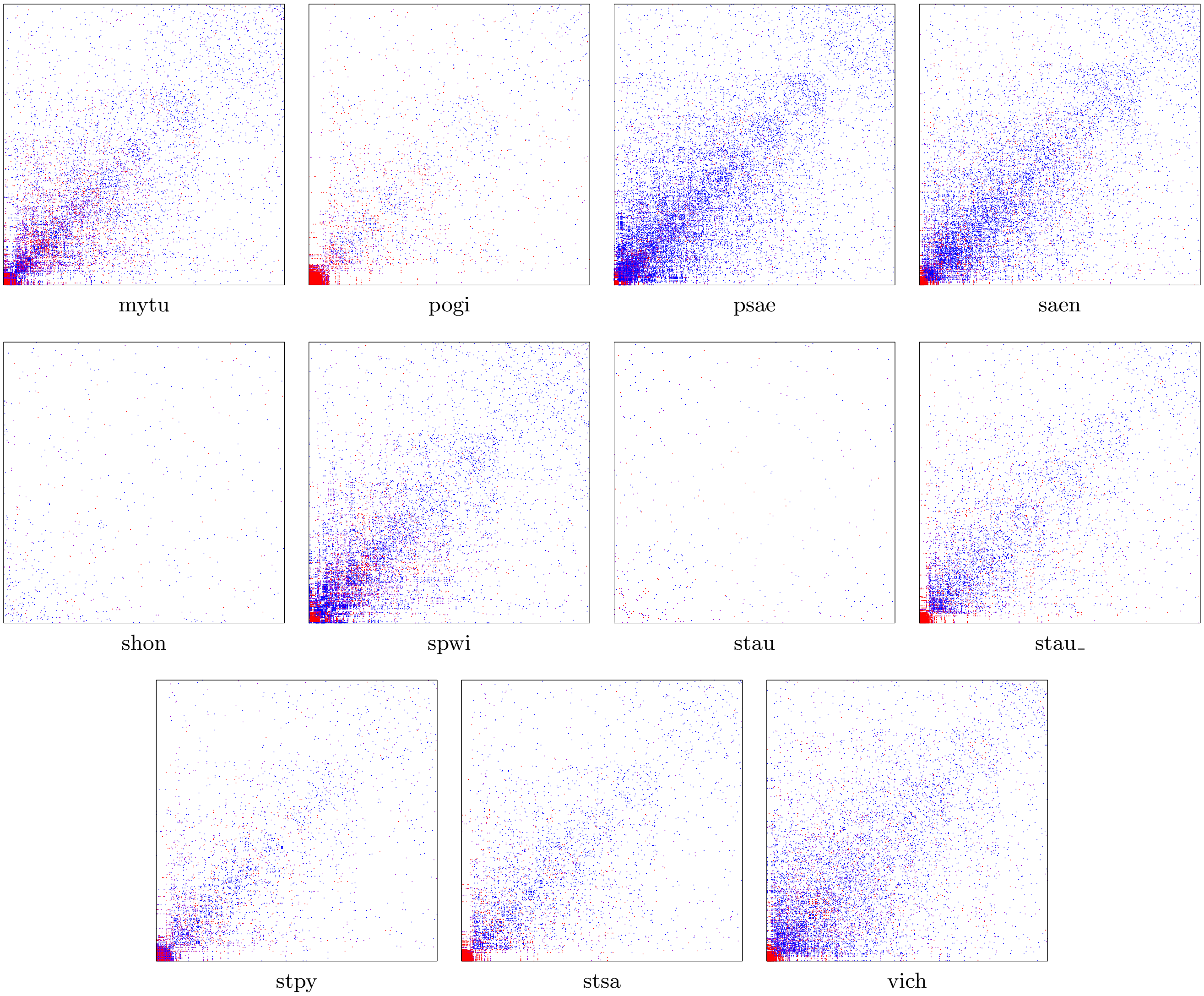


FIG. S7. Adjacency matrices (part 2) of the PPI networks for the DEG-annotated bacterial species reported in Table 1 of the main text. In each matrix, genes are ordered according to the degree of the corresponding protein in the network, in descending order from left to right and from bottom to top. Links among essential genes correspond to red-colored dots, those among nonessential (and non-annotated) genes to blue-colored dots, and those between essential and nonessential (plus non-annotated) genes to a violet-colored dot.

| **Organisms** | **Abbr.** | **STRING** |
|---|---|---|
| Acinetobacter baumannii ATCC 19606 | acba | 470 |
| Aeromonas salmonicida A449 | aesa | 382245 |
| Anabaena variabilis ATCC 29413 | anva | 240292 |
| Azotobacter vinelandii DJ, | azvi | 322710 |
| Bacillus amyloliquefaciens subsp. plantarum | baam | 326423 |
| Bacillus cereus | bace | 1396 |
| Bacillus licheniformis ATCC 14580 | bali | 279010 |
| Bifidobacterium longum | bilo | 216816 |
| Bordetella bronchiseptica | bobr | 257310 |
| Bordetella pertussis 18323 | bope | 568706 |
| Bradyrhizobium japonicum USDA 110 | brja | 224911 |
| Brucella suis 1330 | brsu | 224914 |
| Buchnera aphidicola str, Bp | buap | 261317 |
| Chlamydia muridarum Nigg | chmu | 243161 |
| Clostridium botulinum A str. ATCC 19397 | clbo | 1415774 |
| Clostridium perfringens ATCC 13124 | clpe | 195103 |
| Coxiella burnetii RSA 493 | cobu | 227377 |
| Corynebacterium efficiens YS-314 | coef | 196164 |
| Corynebacterium glutamicum ATCC 13032 | cogl | 196627 |
| Haemophilus ducreyi 35000HP | hadu | 233412 |
| Helicobacter hepaticus ATCC 51449 | hehe | 235279 |
| Klebsiella pneumoniae MGH 78578 | klpn | 573 |
| Lactococcus lactis subsp. cremoris MG1363 | lala | 416870 |
| Leadbetterella byssophila DSM 17132 | leby | 649349 |
| Mycoplasma gallisepticum VA94_7994-1-7P | myga | 708616 |
| Mycobacterium gilvum PYR-GCK | mygi | 350054 |
| Niabella aurantiaca DSM 17617 | niau | 1122605 |
| Nocardia aobensis NBRC 100429 | noao | 1206720 |
| Pseudomonas entomophila L48 | psen | 384676 |
| Pseudomonas putida F1 | pspu | 351746 |
| Pseudomonas syringae pv. phaseolicola 1448A | pssy | 264730 |
| Rhizobium leguminosarum 3841 | rhle | 216596 |
| Rhodobacter sphaeroides ATCC 17025 | rhsp | 349102 |
| Saccharibacillus kuerlensis DSM 22868 | saku | 1123226 |
| Shewanella loihica PV4 | shlo | 323850 |
| Staphylococcus carnosus subsp. carnosus TM300 | stca | 396513 |
| Streptococcus mutans UA159 | stmu | 210007 |
| Streptococcus pneumoniae D39 | stpn | 373153 |
| Synechococcus sp. WH 5701 | sysp | 69042 |
| Vibrio vulnificus YJ016 | vivu | 672 |
| Yersinia enterocolitica subsp. enterocolitica 8081 | yeen | 393305 |
| Yersinia pestis CO92 | yepe | 214092 |

TABLE I. Summary of the selected bacterial dataset. Organism name, abbreviation, STRING code.
